# Source tracking and global distribution of the mobilized tigecycline resistant gene *tet*(X)

**DOI:** 10.1101/2021.07.29.454411

**Authors:** Rong-min Zhang, Jian Sun, Ruan-yang Sun, Min-ge Wang, Chao-yue Cui, Liang-xing Fang, Mei-na Liao, Xiao-qing Lu, Yong-xin Liu, Xiao-Ping Liao, Ya-Hong Liu

## Abstract

The emergence of *tet*(X) genes has compromised the clinical use of the last-line antibiotic tigecycline. We identified 322 (1.21%) *tet*(X) positive samples from 12,829 human microbiome samples distributed in four continents (Asia, Europe, North America and South America) using retrospective data from worldwide. These *tet*(X) genes were dominated by *tet*(X2)-like orthologs but we also identified 12 samples carrying novel *tet*(X) genes, designed *tet*(X15) and *tet*(X16), that were resistant to tigecycline. The metagenomic analysis revealed these *tet*(X) genes distributed in anaerobes dominated by *Bacteroidaceae* (78.89%) of human-gut origin. The transmission of these *tet*(X2)-like orthologs between *Bacteroidaceae* and *Riemerella anatipestifer* was primarily promoted by the mobile elements IS*Bf11* and IS*4351. tet*(X2)-like orthologs was also developed during transmission by mutation to high-level tigecycline resistant determinants *tet*(X15) and *tet*(X16). Further tracing these *tet*(X) in single bacterial isolate from public repository indicated that *tet*(X) genes were present as early as 1960s in *R. anatipestifer* that was the primary *tet*(X) carrier at early stage (before 2000). The *tet*(X2) and non-*tet*(X2) orthologs were primarily distributed in humans and food animals respectively, and non-*tet*(X2) were dominated by *tet*(X3) and *tet*(X4). Genomic comparison indicated these *tet*(X) genes were likely to be generated during *tet*(X) transmission between *Flavobacteriaceae* and *E. coli*/*Acinetobacter* spp.., and IS*CR2* played a key role in the transmission. These results suggest *R. anatipestifer* was the potential ancestral source of *tet*(X) gene. Additionally, *Bacteroidaceae* of human-gut origin was an important hidden reservoir and mutational incubator for the mobile *tet*(X) genes that enabled spread to facultative anaerobes and aerobes.

## Introduction

The first generation of tetracycline antibiotics consisted of tetracycline, chlortetracycline and oxytetracycline and were put into clinical practice in 1952 (1) while the second generation derivatives doxycycline and minocycline were put into use in 1976 (2). These antibiotics have been incorporated into animal feed to improve growth and feed efficiency (3). However, bacterial resistance to the tetracyclines was observed from the very beginning of their usage. To date, more than 65 specific resistant determinants and 9 MDR efflux pump genes of the root nodulation-division (RND) superfamily have been confirmed including AdeABC, AcrAB-TolC and MexAB-OprM (2). These determinants confer resistance to first and second generation tetracyclines and are widely distributed among 130 Gram-negative and Gram-positive bacteria (2).

A third-generation tetracycline (tigecycline) was approved in the United States in 2005 and its use in the EU and China was authorized in 2006 and 2010, respectively (4, 5). Tigecycline has a robust treatment range and includes bacteria resistant to first and second generation tetracyclines (6), and is a ‘last resort’ antibiotic used to treat severe infections caused by carbapenem- and colistin-resistant pathogens (5). Thus, this antibiotic was classified as a critically important antimicrobial by the World Health Organization and its usage is restricted (7). However as early as 1884, the transferable gene *tet*(X) displaying tigecycline insusceptibility was discovered on an R plasmid from a *B. fragilis* isolate of human origin. This was the earliest occurrence of an antibiotic resistance gene (ARG) that directly inactivated tetracyclines (8). The *tet*(X) gene was only functional under aerobic growth conditions because it is a flavin dependent monooxygenase that requires FAD, NADPH, Mg^2+^ and O_2_ to inactive almost all of the tetracycline class (9). In 2001 the existence of *tet*(X1) and *tet*(X2) were confirmed on a transposon from *Bacteroides thetaiotaomicron* of human origin and shared 61.7 and 99.5 % amino acid identity with *tet*(X), respectively. To date, *tet*(X)/*tet*(X2) genes have already spread to 16 countries/regions covering five continents (Asia, Europe, North America, South America and Africa) (2). A small comfort was that these ARGs displayed low level tigecycline resistance (MIC ≤ 2ug.ml^−1^) (2).

The emergence of plasmid-mediated high-level tigecycline resistance encoded by the *tet*(X3) and *tet*(X4) genes in 2019 posed a severe threat to public health (7, 10). Additionally, 10 more *tet*(X) orthologs have been identified and include *tet*(X5) - *tet*(X14)) (11–14). These orthologs were primarily found in food animals especially swine, including *tet*(X3), *tet*(X4), *tet*(X6) and *tet*(X14) firstly detected in *Acinetobacter baumanii*, *Escherichia coli*, *Myroides phaeus* and *Empedobacter stercoris*, respectively. The *tet*(3.2) and *tet*(X5) gene were identified from an *Empedobacter brevis* isolate of shrimp origin and a *Acinetobacter baumannii* isolate of human origin, respectively. All the *tet*(X7) - *tet*(X13) orthologs were identified directly from gut-derived metagenomic libraries, but their host bacteria were unknown. Epidemiological studies (2, 15, 16) indicated the dissemination of these *tet*(X) orthologs were dominated by *tet*(X3) and *tet*(X4) that were primarily detected from *Acinetobacter* spp. and *E. coli*, respectively. Furthermore, *tet*(X3)/*tet*(X4) samples from humans, animals and meat for consumption revealed a prevalence of 0.3 - 66.7 % and the highest level of 66.7 % was detected from pig caecum samples from abattoirs (7). Compared with the *tet*(X3)/*tet*(X4) in animal isolates (6.9%, 73/1060) (7), lower prevalence from human (0.32%, 4/1250) were observed in a retrospective screening of *tet*(X)-carrying clinical isolates (5).

The *tet*(X) genes have been primarily identified using traditional cultural methods and this imposed limitations on their identification including the loss of uncultured bacteria, low-throughput and long processing times. Although the *tet*(X7) - *tet*(X13) orthologs were found directly from gut-associated samples using metagenomic sequencing, the bacterial hosts, relative abundance and propagative characteristics were absent (14). Nonetheless, public repositories are a promising high-throughput resource for exploring antibiotic resistomes. For instance, a retrospective epidemiological study based on the available public bacterial gene datasets revealed that the food chain was a potential dissemination pathway for *mcr-1* (17). Additionally, a metagenomic screening study based on public metagenome datasets revealed a high detection rate of *tet*(X3) (25.4 %) in poultry samples (18). However, there are few studies that utilize data mining for *tet*(X) in public databases (18, 19). In addition, public data repositories including GenBank are a valuable resource for the exploration of novel bacterial species. For instance, a recent study utilized 9,428 metagenomes to reconstruct 154,723 microbial genome bins that generated 4,390 species level genome bins (SGB) including 77 % of which were not present in public repositories (20). Identification of *tet*(X) genes from these SGBs and tracing their distribution in assembly isolates from public repository may provide a new perspective for source tracking of the global spread of *tet*(X).

In the current study, we utilized these data mining techniques and discovered that *tet*(X) had emerged as early as 1960 and the *Riemerella anatipestifer* was its potential ancestral source. Additionally, *Bacteroidac****e****ae* of human gut origin were a hidden reservoir and mutational incubator for mobile *tet*(X) genes that enabled spread to facultative anaerobes and aerobes.

## Methods

### Collection of microbial genomic sequences from human microbiome in retrospective data

A total of 26,728 metagenomic samples of human-microbiome origin deposited in public repositories were downloaded and reconstructed microbial genomic bins (MGBs) to explore new bacterial species in previous studies (20–22). We removed the duplicative samples from these metagenomic samples according their accession number and found a total of 12,829 non-duplicate metagenomic samples from 31 countries. These samples were reconstructed into 202,265 MGBs in previous studies (20–22) (Table S1 and S2) and the online released data were screened for the presence of all known *tet*(X) orthologs using BLAST using an 80 % identity and 70 % hit length cut-off in current study. The prevalence of *tet*(X) in 31 countries were plotted using R version 3.5.3. Phylogenetic analysis for amino acid sequences of all Tet(X) gene products was constructed using neighbor joining with the default parameters in Mega X Version 10.0.5 (23) and alignments were constructed using ESPript 3 (24).

### Functional identification of Tet(X)s

Tigecycline resistance for these gene products was assessed by synthesis of full-length nucleotide sequences of all detected *tet*(X) genes. *EcoR*I and a *Sal*I sites were then added 5′ and 3′ respectively (Tsingke Biological Technology, Beijing, China). The synthesized *tet*(X) genes were cloned into plasmid vector pBAD24 and transformed into competent *E. coli* JM109 as described in our previous study (25). The transconjugants *E. coli* JM109+pBAD24-*tet*(X4) and *E. coli* JM109+pBAD24, were used as positive and negative controls, respectively, as previously described (25). The minimum inhibitory concentration (MIC) for tetracycline, doxycycline, minocycline, tigecycline, eravacycline and omadacycline were determined by the broth microdilution method in accordance with Clinical and Laboratory Standards Institute (CLSI) guidelines. Tetracycline, doxycycline and minocycline breakpoints were interpreted according to the European Committee on Antimicrobial Susceptibility Testing (EUCAST) guidelines (http://www.eucast.org/clinical_breakpoints). The United States FDA criteria was employed to interpret tigecycline breakpoints for *E. coli* and MIC ≥4 mg L^−1^ was considered non-susceptible while eravacycline and omadacycline were uninterpreted with no breakpoint. *E. coli* ATCC 25922 was used as the quality control strain.

### Taxonomic assignment and phylogenetic analysis of *tet* (X)s-carrying MGBs

We obtained 322 *tet*(X)-carrying MGBs from three previous studies (20–22). This group included taxonomic assignments for 196 that had been previously annotated (20) while the remaining 126 were annotated using metaWRAP-Annotate-bins module using the MetaWRAP pipeline and default parameters (26). Briefly, the assembly contigs from each *tet*(X)-carrying MGB was taxonomically profiled using Kraken2 (27) and then this entire metagenomic bin could conservatively and accurately estimate the taxonomic profiles (26).

The phylogenetic structure for the *tet*(X)-carrying MGBs were performed using an automatic PhyloPhlAn (3.0) pipeline (20, 28), through which the phylogeny in Figure 1b was built using 400 universal PhyloPhlAn markers with parameter: “--diversity high --accurate --min_num_markers 80.'’ This pipeline integrates diamond (version 0.9.32), mafft (version 7.464) (29), trimal (version 1.4.rev15) (30) and RAxML (version 8.2.12) (31), and the parameters of these software were set as described previously (20). The phylogenetic tress in Figure 1b were plotted using GraPhlan (version 1.1.3) (32).

**Figure 1.**
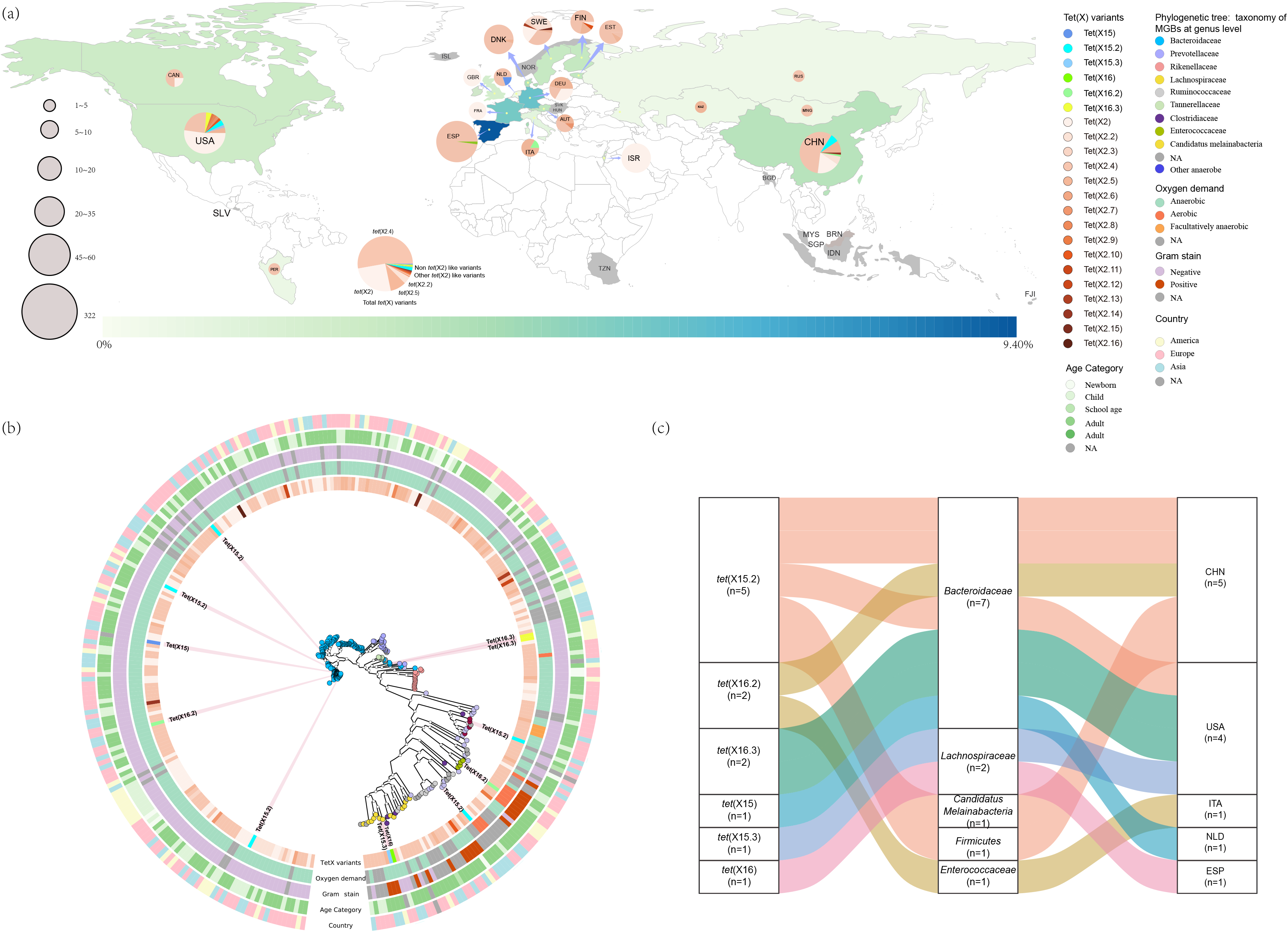
Global distribution of *tet*(X)s from human microbiome. A. World map showed the positive rates of *tet*(X) gene in 19 countries and the colored countries represented the positive rates of *tet*(X) according to the hot map (>0% - 9.40%) at the bottom. The gray countries indicated they were negative for *tet*(X) gene. The size of the pie charts represented the numbers of *tet*(X)-positive MGBs and the colors in the pie charts indicated the composition of *tet*(X) variants. B. PhyloPhlAn analysis of the *tet*(X) carrying MGBs. The taxonomic assignments of the *tet*(X)-carrying MGBs were depicted with colored circles in the phylogenetic tree. The *tet*(X) variants carried by the MGBs, as well as oxygen demand, gram stain, age category and the countries of the *tet*(X)-carrying MGBs were showed in the five colored rings surrounding the phylogenetic tree. C. Distribution of the 12 MGBs carried non-*tet*(X2) genes with tigecycline inactivate function.

### Phylogenetic analyses of the *tet* (X) carrying isolates and evolutionary timescale for the *tet*(X)s from isolates

To further trace the spread of all *tet*(X)s in culturable bacteria isolates, a total of 774,435 bacteria assembled whole genome sequences (WGS) were downloaded from the NCBI database as of November 7^th^, 2020. *tet*(X)-like OFRs were determined using BLATX against all the *tet*(X)s variants mentioned in current study with a minimum similarity of 70% and 100% coverage. The collection date, origin, countries and the bacterial host of the *tet*(X)-positive isolates were retrieved according to their Biosample Number. The phylogenetic structure for the *tet*(X)-carrying isolates were also performed using PhyloPhlAn (3.0) pipeline mentioned above.

To determine the evolutionary history of *tet*(X)s, the earliest emergency of *tet*(X) variants (with collection date) with a 388 amino acid (aa) length were applied to generate a chronogram using Bayesian evolutionary analysis version 1.10 (33). For all model combinations, three independent chains of 100 million generations each were run to ensure convergence with sampling every 1,000 iterations. Tracer v1.7.1 was used to assess convergence using all parameter effective sampling sizes of > 200 (34). LogCombiner v2.6.1 was used to combine tree files and a maximum clade credibility tree was created using TreeAnnotator v2.6.0 (34). Tree annotation was visualized using iTOL (35) and FigTree version 1.4.2.

### Annotation and comparison of the genomic region flanking the *tet*(X) gene

The *tet*(X)-carrying contig were extracted from the MGBs of metagenomic analysis and isolates from public repository. CD-HIT was employed to group *tet*(X)-carrying full length contigs using a cutoff with a minimum similarity of 97 % over 97 % of the query coverage (36). These *tet*(X)-carrying contigs were annotated using Prokka (37) and in conjunction with standalone BLAST analyses against the ResFinder (38) and ISfinder (39) databases to cross-validate ARGs and mobile genetic elements, respectively.

## Results

### Identification of Tet(X) orthologs

We accessed a total of 202,265 MGBs constructed from 12,829 metagenomic samples of human-microbiome origin and found 322 (1.21 %) of them encoded 388 aa proteins with 96.13 - 100% similarity with Tet(X) orthologs reported in a previous study (11) (Table S2). In particular, 96.27 % (310/322) of them shared 98.71 - 100 % identity with Tet(X2) (Acc. No. AJ311171) and phylogenetically could be grouped into 15 *tet*(X2)-like clades (Figure S1 and S2, Table S3). The remining 3.73 % (12/322) shared < 98.20 % identical with the known Tet(X). Since there was not a criterion for assignment of Tet(X) orthologs in previous studies (11, 14), we temporarily designated these *tet*(X2)-like orthologs as *tet*(X2.2) to *tet*(X2.16). These orthologs represented a large proportion of samples (n=12,829) from 19 countries and were therefore likely represented a relatively comprehensive assessment of *tet*(X2)-like orthologs. We therefore tentatively used the lowest cutoff of 98.20 % between *tet*(X2)-like orthologs (*tet*(X2.15) *vs tet*(X2.16)) for assignment of novel *tet*(X) orthologs. Based on this, we found two new *tet*(X) orthologs and their subtypes shared less than 98.20% amino acid identity with their closest neighbors and most resembled *tet*(X) orthologs in the phylogeny designed *tet*(X15), *tet*(X15.2), *tet*(X15.3), *tet*(X16), *tet*(X16.2) and *tet*(X16.3) (Figure S1, Table S3). Most of these *tet*(X) orthologs found from metagenomic analysis were not present in the NCBI database with the exceptions of *tet*(X2) (Acc. No. AJ311171), *tet*(X2.4) (Acc. No. JQ990987) and *tet*(X16.2) (Acc. No. KU547718.1) (Table S3).

### Resistance phenotypes of *tet*(X) orthologs

Both the *tet*(X15) and *tet*(X16) groups found from metagenomic analysis of human-gut origin (Figure 1c) in the *E. coli* JM109 were resistant to tigecycline (MICs 8 to 16 ml/liter), and exhibited high MIC to the fourth-generation antibiotics omadacycline (MICs 16 to 32 ml/liter) and eravacycline (MICs 2 to 4 ml/liter) (Table S4). In addition, all the non-*tet*(X2) exhibited resistance to tetracycline (MICs 128 to 256 ml/liter), doxycycline (MICs 32 to 64 ml/liter) and minocycline (MICs 16 to 32 ml/liter) (Table S4). All the *tet*(X2) like orthologs were susceptible to tigecycline (MICs 0.25 to 1 ml/liter) but resistant to tetracycline (MICs 8 to 256/liter). Almost all of them were resistant to doxycycline (MICs 8 to 64 ml/liter) and minocycline (MICs 2 to 32 ml/liter), excluding *tet*(X2.10) that intermediate to doxycycline (1 ml/liter) and minocycline (MICs 0.5 ml/liter). Most of these *tet*(X2) like orthologs exhibited MIC of omadacycline from 0.5 to 4 ml/liter, and eravacycline from 0.0125 to 1 ml/liter (Table S4). Three *tet*(X2) like orthologs *tet*(X2), *tet*(X2.4) and *tet*(X2.15) showed relatively higher MICs to omadacycline (8 to 16 ml/liter) and eravacycline (1 to 4 ml/liter).

### Global distribution and taxonomic assignment of *tet*(X) carrying MGBs

The 322 *tet*(X)s carrying MGBs were detected in 19 countries including Europe (n=12), Asia (n=4) and America (n=3) but were absent in Oceanica and Africa (Figure 1a, Table S5). The prevalence of *tet*(X) in European countries (0 - 9.40 %) was more complex than Asian (0 - 3.17 %) and American countries (0 - 2.44 %) although the positive rates for *tet*(X) in these three continents were not significantly (P>0.05) different (Figure S3). The prevalence of *tet*(X) was highest in Spain (9.40 %), followed by Germany (5.14 %), France (3.82 %), Denmark (3.34 %) and China (3.17 %) and the remaining countries were < 3 % (Figure 1a and Table S5).

The bacterial taxonomy assignment indicated that all the 322 *tet*(X)-carrying MGB could be assigned to five phyla and was dominated by *Bacteroidetes* (78.89 %, 254/322) followed by *Firmicutes* (16.78 %, 47/322), *Proteobacteria* (1.24 %, 4/322), *Candidatus Melainabacteria* (1.24%, 4/322) and *Fusobacteria* (0.62%, 2/322) (Figure 1b). Furthermore, 96.68% (301/322) were classified to the family level and most of them were also belonging to *Bacteroidaceae* (70.10%, 211/301) (Figure 1b and Table S6). Furthermore, 72.30% (154/211) of these *tet*(X) carrying *Bacteroidaceae* MGBs could be further divided into 14 *Bacteroides* species that were dominated by *Bacteroides vulgatus* (37.67 %, 58/154), *Bacteroides uniformis* (20.78 %, 32/154), *Bacteroides dorei* (14.94 %, 23/154), *Bacteroides ovatus* (5.84 %, 9/154), *Bacteroides fragilis* (3.90 %, 6/154) and *Bacteroides caccae* (3.25 %, 5/154) (Table S7).

The *tet*(X2)-like positive MGBs were distributed across these 19 *tet*(X) positive countries. *tet*(X2.4) (55.02 %, 170/309), *tet*(X2) (26.86 %, 83/309), *tet*(X2.5) (9.32 %, 30/309) and *tet*(X2.2) (4.53 %, 14/309) totaled 96.11 % (297/309) and prevailed over other *tet*(X2)-like orthologs (≤ 0.6 %, 2/309) (Figure S4). Only 81.08% (240/296) of these four predominant *tet*(X) carrying MGBs could be assigned at genus level and most of them were also identified in *Bacteroides* (71.67%,172/240), followed by *Prevotella* (6.67%, 16/240) and *Alistipes* (5%, 12/240) (Table S8). Additionally, the non-*tet*(X2) orthologs included 75 % (9/12) that were distributed in China and USA. Of which, *tet*(X15.2) was the most prevalent ortholog (Figure 1c). The remaining three non-*tet*(X2) carrying MGBs were distributed in Europe and the single *tet*(X16.2) ortholog was carried by an *Enterococcaceae* from Italy. Almost all *tet*(X)-carrying MGBs were collected from human stools excluding only one *tet*(X2.4)-carrying MGB identified as an *Enterococcus* spp. from an oral cavity sample (Table S8).

The 322 *tet*(X) carrying MGBs included 196 from the 4,390 MGBs and their average abundance in human microbiomes has been calculated in previous study (Table S8) (20). The average abundance of these 196 *tet*(X)-positive MGBs was 5.97 ± 3.89 and significantly higher than that of the total 4,390 MGBs (1.76 ± 3.74) (20).

### Culturable isolates from public repository insight into the distribution and evolutionary timescale of Tet(X)s

To further trace the distribution of these *tet*(X)s in culturable isolates, we examined 774,435 WGS of bacterial isolates present in GenBank and only 0.12 % (896/774,435) carried an ORF with > 70 % amino acid identity with the known *tet*(X) genes including the novel ones found in the current study (Table S9). The PhyloPhlAn analysis indicated that the facultative anaerobe clade was phylogenetically distinct between anaerobes and aerobes (Figure 2a). These *tet*(X) genes were found in 17 bacterial families that were dominated by aerobes including *Moraxellaceae* (279/896, 31.17 %), *Enterobacteriaceae* (208/896, 23.24 %) and *Weeksellaceae* (16.20 %, 145/896). The *Bacteroidaceae* (20.67 %, 185/896) were also an important anaerobic carrier for *tet*(X2) like orthologs (Figure 2a and Table S9). Three *tet*(X) orthologs were most prevalent and included *tet*(X2)-like (35.71 %, 320/896), *tet*(X3) (27.34 %, 245/896) and *tet*(X4) (21.99 %, 199/896) (Table S9). Interestingly, different bacterial families from variant hosts were preference for carrying specific *tet*(X) ortholog (Figure 2a). Almost all the *Bacteroidaceae* (98.91 %, 182/184) and most *Weeksellaceae* (63.45 %, 92/145) isolates carried *tet*(X2)-like orthologs and were primarily from human samples (38.64 %, 114/295). Additionally, all *tet*(X3) were detected from *Acinetobacter* spp. and almost all *tet*(X4) (88.89 %, 176/197) were carried by *E. coli* and these were primarily from food animals (66.59 %, 295/443) including pigs, chickens, ducks, cattle and geese.

**Figure 2.**
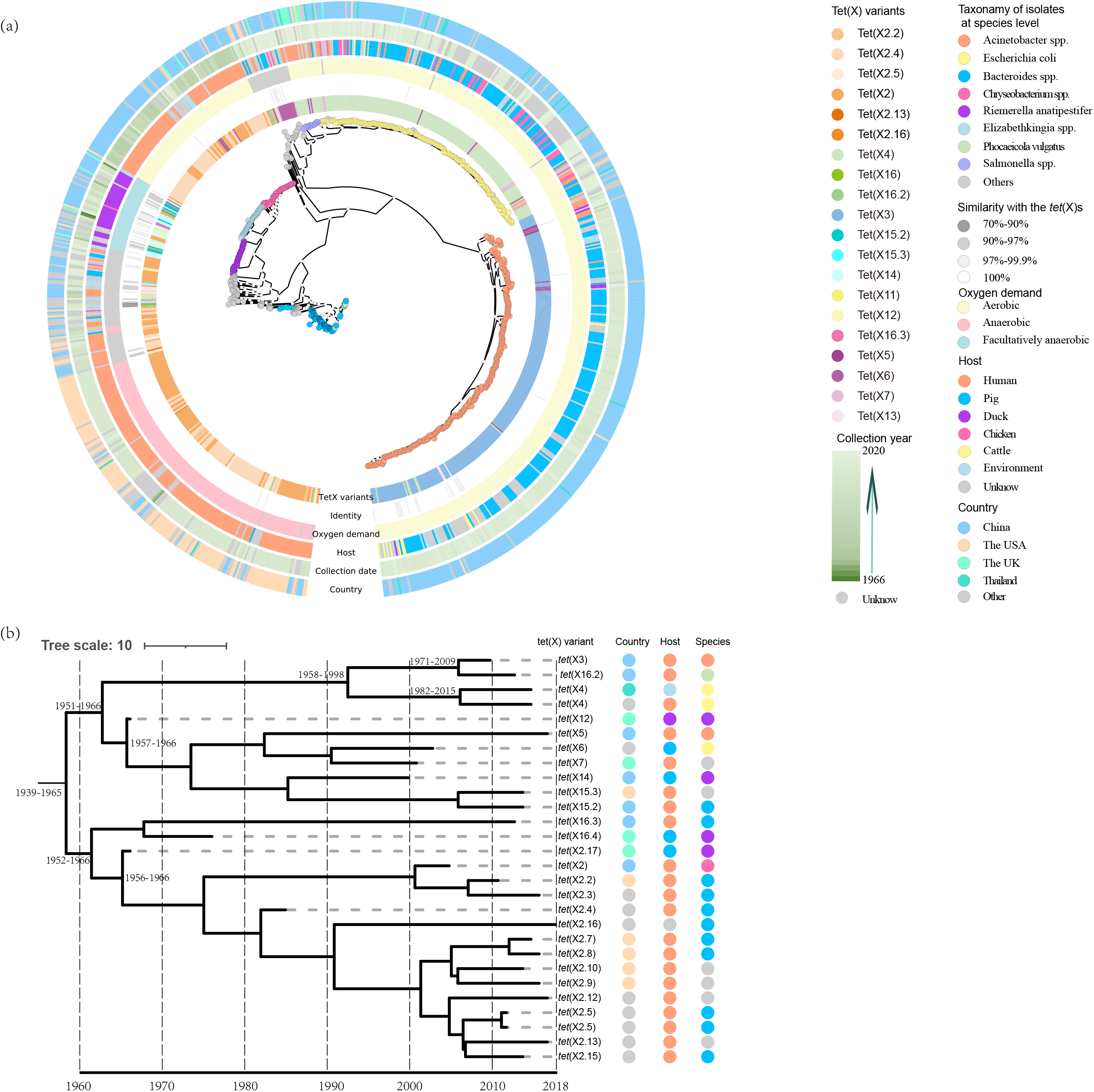
Culturable isolates insight into *tet*(X) distribution patterns. A. PhyloPhlAn analysis of the *tet*(X)-carrying isolates from the public repository. The species of the *tet*(X)-carrying isolates were depicted with colored circles in the phylogenetic tree. The information of the *tet*(X) carrying isolates including *tet*(X) variants, oxygen demand, host, collection date and country were showed in the six colored rings surrounding the phylogenetic tree. B. Dates of lineage divergence of the earliest *tet*(X) orthologs as determined using Bayesian phylogenetic inference. The *tet*(X) variants, countries, host and species of these isolates were shown at the right region.

The *tet*(X2)-like orthologs were detected prior to 8.41 ± 6.17 years ago and earlier than *tet*(X3) (4.00 ± 1.12) and *tet*(X4) (2.38 ± 1.34). To be noted, the collection dates of three *R. anatipestifer* isolates from UK duck samples were prior to 1980. One isolate (BioSample: SAMN09912225) was collected in 1966 and was positive for *tet*(X12) and a *tet*(X2)-like gene that only differed in a single amino acid from *tet*(X2) was designed *tet*(X2.17). The two other isolates were collected in 1976. One (BioSample: SAMN09912224) carried a *tet*(X2.17) gene and another (BioSample: SAMN09912221) carried a *tet*(X) gene shared 98.20% similarity with *tet*(X16), designed *tet*(X16.4).

Date for lineage divergence of the earliest occurred *tet*(X) orthologs were produced by Bayesian phylogenetic inference (Figure 2b). The analysis indicated a mean rate of 0.28 SNP per year for these *tet*(X) during 1966-2018. The most-recent common ancestor (MRCA) of all *tet*(X) from this study was approximately from 1939 to1965, and the tracer analysis revealed the presence of *tet*(X) most likely occurred in AD 1953 (95% highest posterior distributor (HPD) AD). Two main lineages that originated from *tet*(X2.17) and *tet*(X12), respectively, were observed in this phylogeny and both *tet*(X) orthologs were collected from the *Riemerella anatipestifer* of duck origin from the UK in 1966 year.

### Annotation and comparison of the *tet*(X) genomic environment

A total of 1218 *tet*(X)-carrying contigs ranging from 1,190 to 931,600 bp were retrieved from the metagenome and bacterial-isolate data. These contigs were grouped into 455 clusters that carried a range of 1 - 48 contigs (Table S10). In each cluster, longer contig shared more than 97% coverage and more than 97% similarity with shorter contig. The high coverage and similarity of these contigs indicated that these *tet*(X) could spread among each cluster (Table 10). This indicated *tet*(X) orthologs could spread among a great diversity of host including human, animal and environment (Figure 3). Both humans and pigs were the primary *tet*(X) hosts. *tet*(X2)-like *tet*(X3), *tet*(X4), *tet*(X7) and *tet*(X16.3) have been found in humans as well as *tet*(X2)-like, *tet*(X3), *tet*(X4) and *tet*(X6) in pigs. However, only *tet*(X2)-like and *tet*(X3) orthologs could transfer between these two hosts (Figure 3a). Interestingly, *tet*(X2)-like orthologs could hitch a great diversity of vehicles to spread between humans and pigs and these included *Bacteroides* spp., *R. anatipestifer* and *Chryseobacterium* spp.. The remaining *tet*(X) genes were spread only *via* special species between different hosts. For instance, the *tet*(X3) gene could only be transited by *Acinetobacter spp*. and spread between pigs and other hosts including pigeons, cattle, geese, ducks and humans (Figure 3a). Additionally, the *tet*(X4) in the genomic array *rdm*C-*tet*(X4)-*Δ*IS*CR2* could spread among wild birds, humans, pigs, chickens and the environment (Figure 3b).

**Figure 3.**
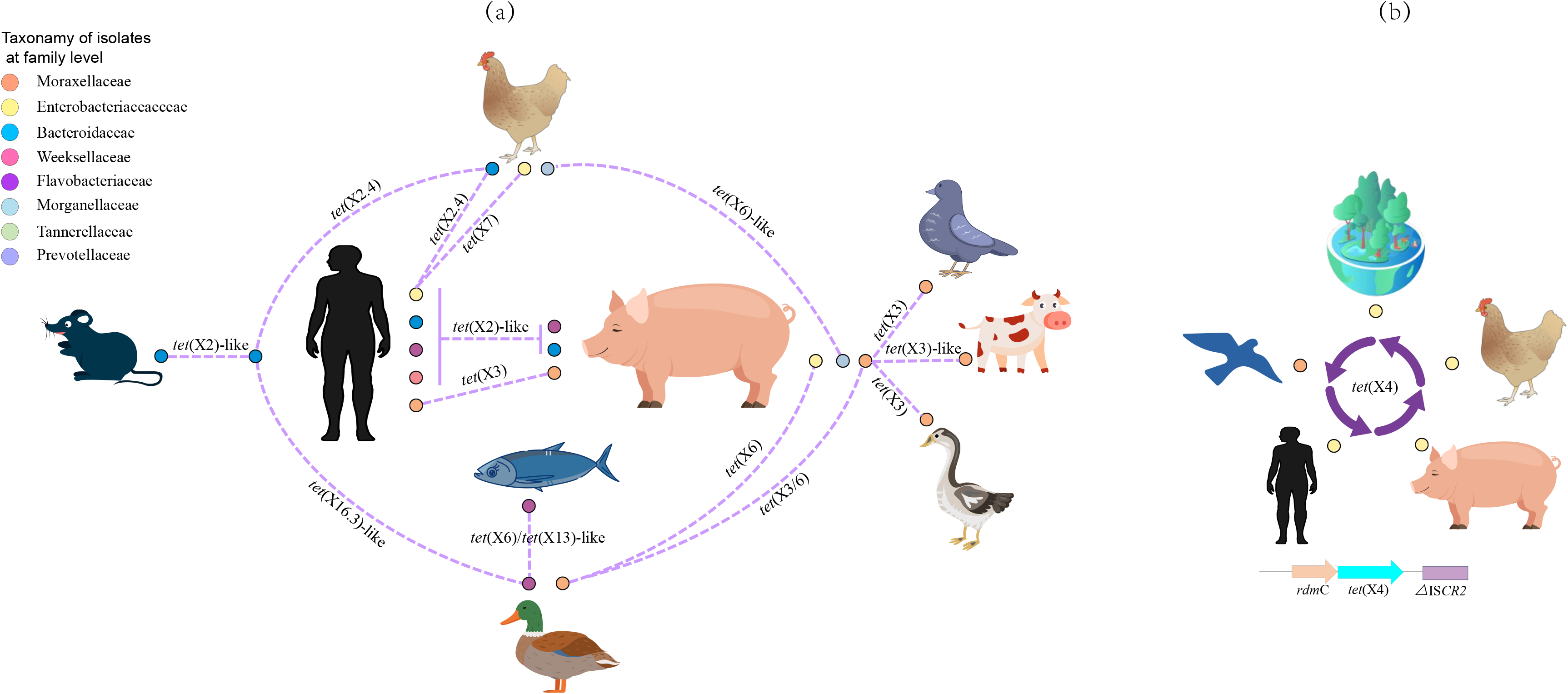
Possible transmission routes of the *tet*(X) genes. The colored circles surrounding the hosts represented the *tet*(X) carrying bacterial families. The dotted lines represented the possible transmission routes of the *tet*(X) genes between different hosts.

Genomic annotation and comparisons indicated that other ARGs frequently flanked the *tet*(X) gene and these included *erm*F, *aad*K, *tet*(Q) and *bla*_OXA347_ that conferred resistance to erythrocin, streptomycin, tetracycline and ampicillin, respectively (Figure 4a and 4b). Of which, the *erm*(F) gene was the most frequently flanking the *tet*(X) gene (n=152) and their genomic environment were clustered into two types according to their relative position: *erm*F located upstream (15.79 %, 24/152) and downstream (80.92 %, 123/152) of *tet*(X) gene (Figure 4a and 4b). The upstream *erm*F always formed a conserved structure *tnpF*-*erm*F-*tet*(X1)-*tet*(X2)/*tet*(X2.2)-*aad*K (n=24) (Figure 4a). This structure was also present in a conjugative transposon CTnDOT of *Bacteroides* origin (40), but the *aad*K (930 bp) was replaced by another aminoglycoside ARG *aad*S (903 bp) in CTnDOT and these two ARGs shared a 96.77 % identity at the nucleotide level. Interestingly, the *tet*(X) orthologs and their genomic contexts were more diverse when *erm*F was located immediately downstream of *tet*(X). These *tet*(X) genes included 12 orthologs that were dominated by *tet*(X2.4) and *tet*(X2.5) (Figure 4b, S1 text A1). In addition, these *tet*(X) genes were able to spread among anaerobes, aerobes and facultative anaerobes (text S1).

**Figure 4.**
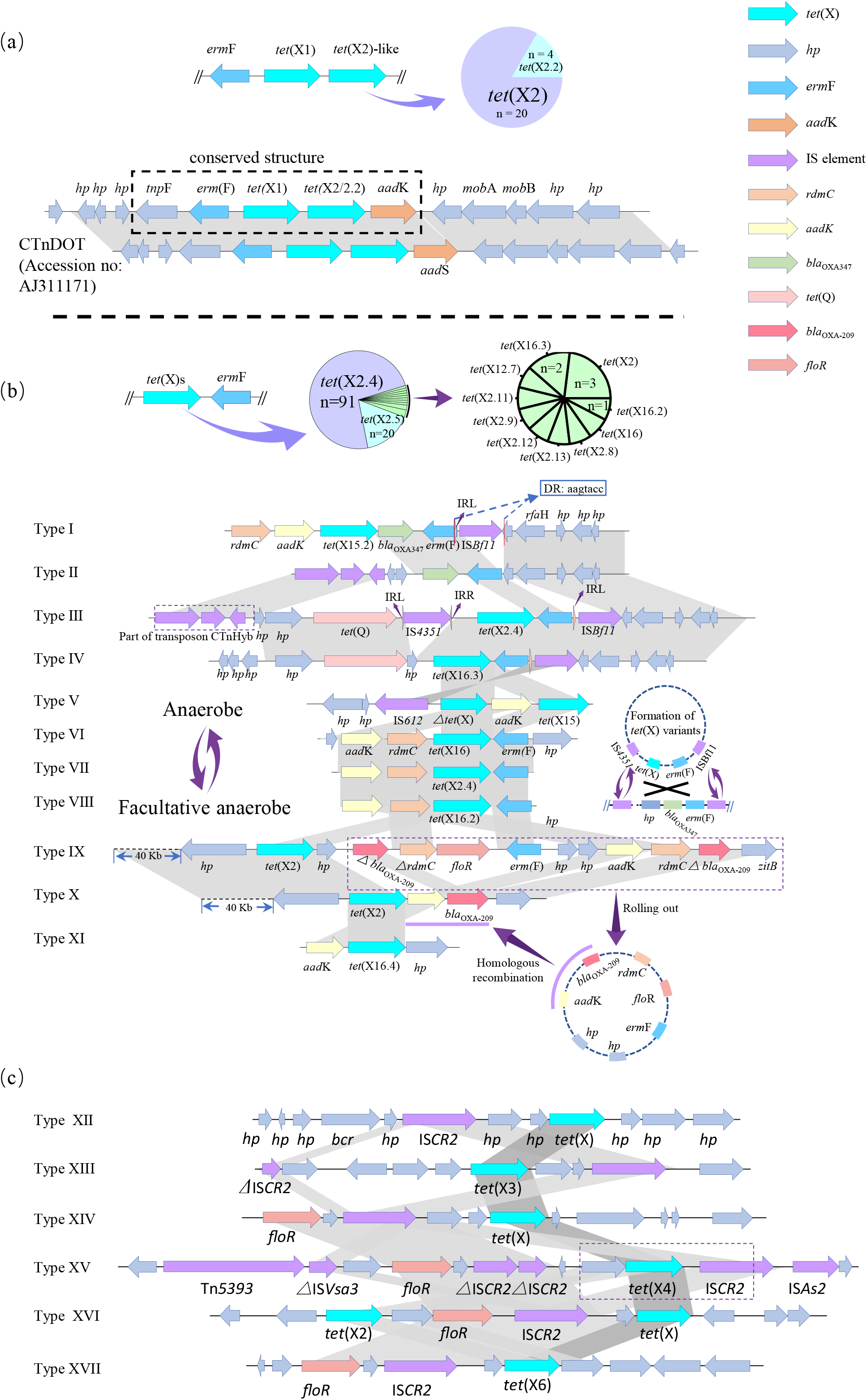
Comparison of *tet*(X) genomic environment. A. The genomic comparison of *erm*(F) gene located upstream of *tet*(X2)-like genes. The proportions of the *tet*(X2) and *tet*(X2.2) located downstream *tet*(X1) were showed in the pie chart. B. The genomic comparison of *tet*(X) genes located downstream *erm*(F). The proportions of the *tet*(X) variants located downstream of *erm*(F) were showed in the pie chart. The possible mechanisms of non-*tet*(X2) formations were showed in the two circles plotted with dotted line. C. Genomic comparison of the regions flanking *tet*(X3) and *tet*(X4) among *Flavobacteriaceae*, *Acinetobacter* and *E. coli*. Arrows indicate the directions of transcription of the genes, and different genes are shown in different colors. Regions of ≥ 99.0% nucleotide sequence identity are shaded light grey. Regions of 77% - 91% nucleotide sequence identity are shaded dark grey. The *Δ* symbol indicates a truncated gene. IS, insertion sequence. See Table S11 for genomic Type I - XVII definitions.

## Discussion and conclusion

### Source tracking of the *tet*(X) orthologs

One of the most significant findings in current study was that *tet*(X) emerged as early as 1960s in *R. anatipestifer* of duck origin, which was earlier than that had been previously reported in 1980s (8) but we found *tet*(X) was present. A previous epidemiological study indicated the MRCA of these *tet*(X)s orthologs likely occurred 9900 years ago (7887 BC) (15). However, we also estimated the MRCA of these *tet*(X) genes but determined it occurred only 68 years ago (1953 AD) that was only one year following the introduction of tetracycline in clinics. This was consistent with the streptomycin, erythrocin and florfenicol, in which their resistance also emerged over the ensuing years following their introduction in clinic [4]. In this analysis, we screened for the presence of *tet*(X) orthologs from public repositories on a large scale (774,435 isolates) and the earliest emergence of 28 *tet*(X) variants was selected to perform MRCA estimations (Figure 2b). The time of the earliest occurrences of *tet*(X) were an important factor for the chronogram phylogeny construction using the Bayesian evolutionary analysis. Thus, these data should be validated to confirm these results.

The *Flavobacteriaceae* have been recognized as a potential ancestral source of the tigecycline resistance gene *tet*(X) (5). We found that both the two earliest (1966) occurring *tet*(X) orthologs (*tet*(X2.17) and *tet*(X12)) harbored by *R. anatipestifer* that is also a bacterial species member belonging to *Flavobacteriaceae* family. Meanwhile, before 2000 only a total of seven isolates were confirmed as *tet*(X) gene carrier and five of them were also identified as *R. anatipestifer* (Table S9). Among the 896 *tet*(X)-carrying isolates from public repository, 71 *tet*(X)-carrying were *Flavobacteriaceae* including *R. anatipestifer* that accounted for a large number (64.79%, 46/71) versus all other *Flavobacteriaceae* members (Figure 2a). Additionally, *R. anatipestifer* harbored a great diversity of *tet*(X) variants and only 23.94 % (17/71) carried the known *tet*(X)s including *tet*(X12), *tet*(X14), *tet*(X2.17) and *tet*(X16.4) (Figure 2a). The remaining carried other *tet*(X) orthologs that shared 94.9 - 99.7 % similarity with their most closely related *tet*(X) ortholog (Table S9). A recent study indicated the poultry pathogen *R. anatipestifer* appears to be a reservoir for Tet(X) tigecycline resistance (41).These indicated that *R. anatipestifer* was most likely the ancestral source of the tigecycline resistance gene *tet*(X).

A great diversity for *tet*(X) and their flaking genomic contexts was observed when the *erm*F gene was present downstream of *tet*(X) (Figure 4b and text S1). A comparison of the *tet*(X) genomic contexts from MGBs and culturable isolates yielded 11 genomic backbones that were associated with the formation of non-*tet*(X2) orthologs found in the current study (Figure 4b). The non-*tet*(X2) orthologs were likely to be generated from *tet*(X2)-like orthologs during their transmission between anaerobes or between anaerobes and facultative anaerobes (Figure 4b and text S1). IS*Bf11* and IS*4351* played important roles in their transmission between anaerobes that was dominated by *Bacteroides* spp. where a mobile cyclic structure was speculated based on genomic Types I - IV (Figure 4b and text S1), and the *tet*(X15)-like and *tet*(X16)-like genes were likely to be generated during the transmission of this mobile structure. In addition, a self-transmissible cyclic structure lacking IS elements was also hypothesized based on the genomic Types V - XI. This structure likely transferred between *Bacteroides* spp. and *R. anatipestifer* and *tet*(X16.4) was most likely generated during the transmission. In addition, genomic comparisons indicated that these *tet*(X) genes were also able to spread between *Flavobacteriaceae* and *E. coli* as well as between *Flavobacteriaceae* and *Acinetobacter* sp. (Figure 4c and text S1). We have demonstrated that *R. anatipestifer*, a *Flavobacteriaceae* family member, was a potential ancestral source of *tet*(X) and the new *tet*(X) orthologs were likely to be produced during their transmission. The high similarity of the nucleotide sequences flanking *tet*(X3) and *tet*(X4) (Figure 4c and text S1) suggested that these two genes were also derived from *Flavobacteriaceae* and IS*CR2* played a key role in this process (Figure 4c and text S1).

### Global distribution of *tet*(X) orthologs

The human microbiome plays an important role in public health. Here, we first determine the *tet*(X) prevalence in the human microbiome using a large-scale survey of 12,829 samples (20–22). A total of 16 *tet*(X2)-like and two new non-*tet*(X2) orthologs have been identified directly in the human stool samples. Since there was not standard for assignment of the new found *tet*(X) orthologs, and it was necessary to distinguish the different *tet*(X) orthologs in current study. Thus, we temporarily set up a criterion for the assignment of *tet*(X) orthologs. This maybe not comprehensive as the assignment for mobile colistin resistance (*mcr*) genes (42) which have established a platform in NCBI to confirm and allocate the allele numbers for new *mcr* gene. An allele numbers assignment for *tet*(X)s should be established urgently.

We found a prevalence for *tet*(X) at 1.21 % (322/26,548) that was higher than for *E. coli* and *K. pneumoniae* from hospital isolates (0.32%, 4/1520) (5) indicating that traditional culture methods have underestimated the prevalence of *tet*(X). Our results were similar to an epidemiological study that detected the *bla*_NDM_ and *mcr-1* genes directly from samples that was higher than for the *E. coli* isolates (43).

The *tet*(X2)-like genes carrier in human microbiomes were dominated by the *Bacteroidaceae* in contrast to previous epidemiological studies where *tet*(X3) and *tet*(X4) were primarily carried by *A. baumannii* and *E. coli*, respectively (2, 15). The *Bacteroides* are predominant anaerobes estimated to account for 25 - 30% of human gut microflora (44) while the *Enterobacteriaceae* normally constitutes only 0.1 - 1 % (45). We also found that the average abundance of *tet*(X)-carrying MGBs (5.97 ± 3.89) annotated as *Bacteroidaceae* prevailed over species-level genome bins (1.76 ± 3.74) in the human microbiome. This was likely the reason for the absence of *tet*(X3) and *tet*(X4) in our human microbiome analyses. Although *tet*(X) genes are inactive in anaerobes, the high abundance of *tet*(X2)-carrying MGBs and a variety of non-*tet*(X2)-like orthologs found in the current study indicated that the *Bacteroidaceae* were an important reservoir and mutational incubator for the mobile *tet*(X) orthologs in the human microbiome (Figure 5). Furthermore, the *Bacteroidaceae* could generate new non-*tet*(X2) orthologs with tigecycline inactivation functions, and the comparison of the *tet*(X) genomic environment suggested that these non-*tet*(X2) enabled transfer to facultative anaerobes and aerobes (Figure 5).

**Figure 5.**
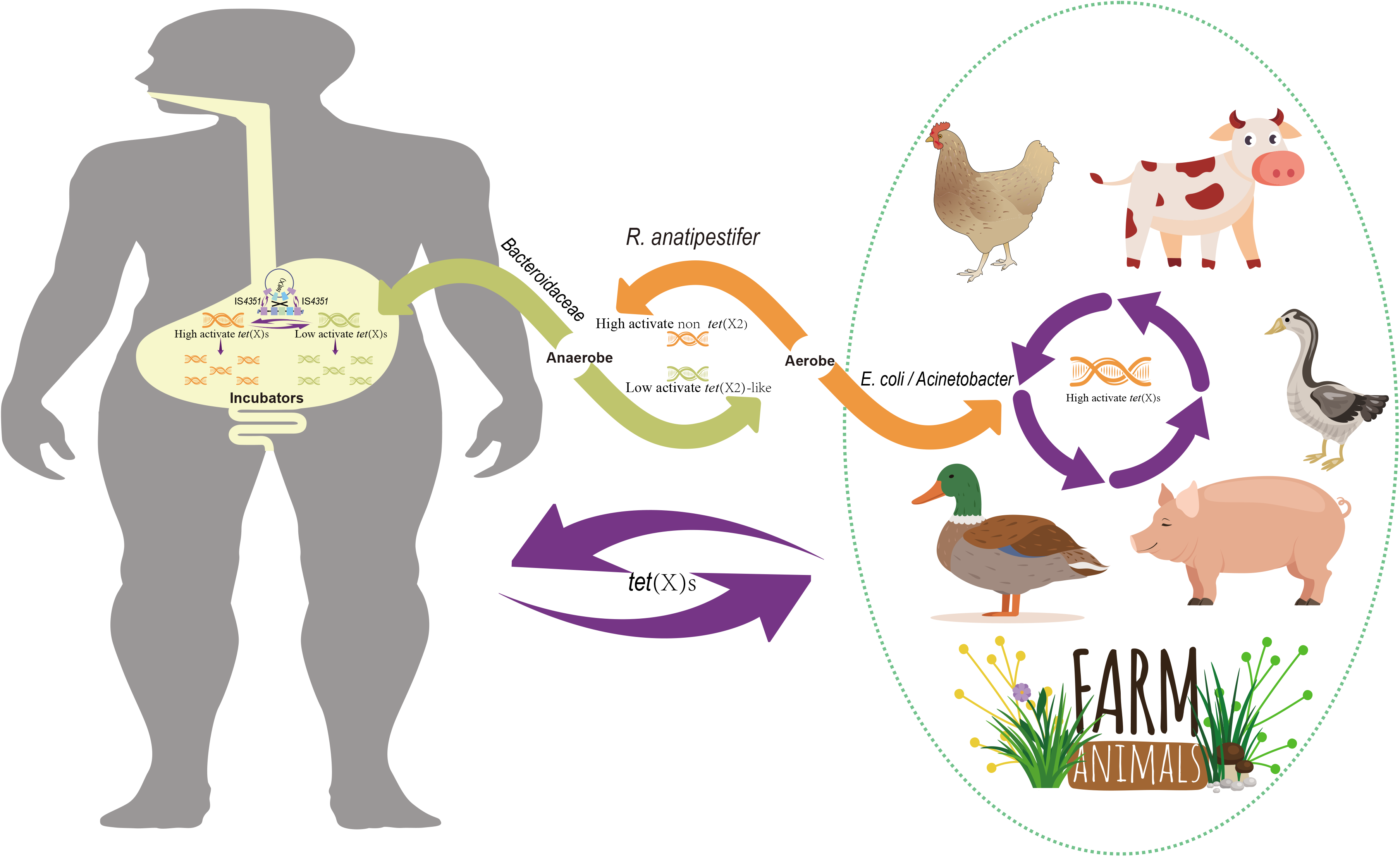
Potential origin and main transmission routes of the *tet*(X) genes.

We have demonstrated *R. anatipestifer* as potential ancestral source of *tet*(X) genes, and the earliest emergence of high-level tigecycline resistance genes *tet*(X3) and *tet*(X4) were likely to be 1971 and 1982, respectively, both of which were earlier than the clinical introduction of tigecycline in 2005. This was further evidence that the use of even older antibiotic tetracycline may contribute to the resistance to newer antibiotics (4). The *tet*(X) distributions from culturable isolates indicated that *tet*(X2)-like and non-*tet*(X2) orthologs were prevalent in anaerobes and aerobes respectively (Figure 5). Since Tet(X) is active only in an aerobic environment, the non-*tet*(X2)-like orthologs with tigecycline inactivate function tended to be captured by aerobes under tetracycline selective pressure (Figure 5). This was likely to be the reason that non *tet*(X2) orthologs primarily distributed in aerobes but the high prevalence of *tet*(X2)-like orthologs in *Bacteroidaceae* from human microbiome need to be further explored.

Taxonomic assignments for *tet*(X)-carrying MGBs were estimated using bioinformatic methods that may not be as precise as cultural methods but such an approach has proved feasible (20, 46). Another limitation of this approach was challenging to combine the chromosome with their respective plasmid sequences (47). Therefore, these *tet*(X)-carrying contexts from MGBs in the current study were likely to be chromosome-borne. This differed with *tet*(X3)/*tet*(X4) that are present in a variety of plasmids and IS*CR2* was an essential element for their mobilization (2, 15). Transmission between *Bacteroidaceae* of these *tet*(X)s orthologs was primarily mediated by the CTnDOT-like conjugative transposon and *erm*F-related IS elements including IS*Bf11* and IS*4351*. CTnDOT has been reported to harbor *erm*F, *tet*(X1) and *tet*(X2) in *Bacteroides* (40) and conjugative transposons can also insert into co-resident plasmids in addition to the chromosome (48). Therefore, conjugative transposons have been found in numerous genera including *Enterococcus*, *Streptococcus*., *Lactococcus*, *Butyrivibrio*, *Clostridium*, *Salmonella*, *Pseudomonas*, *Mezorhizobium* and *Vibrio* (40). The erythromycin resistance *erm*F gene is frequently reported in *R. anatipestifer* and *Bacteroides* spp. isolates (49, 50). The IS *Bf11* and IS*4351* flanking *tet*(X) in the Type III genomic contexts have also been previously identified (51) and reveals that this mobile structure has spread to China, the USA, France, Denmark, Sweden and Belgium. In addition, the IS*CR2* element belonging to the IS*91* family has been described in the first report of *tet*(X3) and *tet*(X4), both downstream and upstream of *tet*(X3). This IS element could form a mobile amplicon and this was demonstrated using inverse PCR experiments (7) and in an *Acinetobacter towneri* isolate flanking region of *tet*(X6) (52). We also identified this IS element located upstream of *tet*(X6) and indicated that this IS element can play an important role in transmission of non-*tet*(X2) orthologs with tigecycline inactivation functions in the *Flavobacteriaceae, E. coli* and *Acinetobacter*.

In conclusion, we conducted an analysis integrated human gut metagenome and global bacterial isolates to trace the origin and distribution of *tet*(X) gene. The *tet*(X) gene emerged as early as 1960 and the *R. anatipestifer* was an ancestral source of *tet*(X). The *tet*(X3)-carrying *Acinetobacter* spp. and *tet*(X4)-carrying *E. coli* were prevalent in food animals and these two *tet*(X)s were likely formed during the transmission of *tet*(X)s between *Flavobacteriaceae* and *E. coli*/*Acinetobacter*. The IS*CR2* element played a key role in the transmission. The *tet*(X2)-like orthologs enriched in the anaerobes that was dominated by *Bacteroidaceae* of human-gut origin and could transfer between these anaerobes. The mobile elements CTnDOT, IS*Bf11* and IS*4351* played important roles in the transmission. The low-level tigecycline resistance *tet*(X2)-like gene could mutate to high-level tigecycline resistant determinants that could spread to facultative anaerobes and aerobes. *Bacteroidaceae* present in the human gut were an important reservoir and mutational incubator for *tet*(X) genes.

## Acknowledgments

This work was supported by the National Natural Science Foundation of China (31730097), the Program for Changjiang Scholars and Innovative Research Team in University of Ministry of Education of China IRT_17R39, the Local Innovative and Research Teams Project of Guangdong Pearl River Talents Program (2019BT02N054), and the 111 Project (D20008).

The authors are grateful to the previous studies that provide a large-scale of microbial genomic bins for the construction of the metagenomic analysis in current study.

## Supplementary files

Text S1. Description of the comparison of *tet*(X) genomic-environment in Figure 4 and S5.

Figure S1. Phylogenetic analysis of the Tet(X) at amino acid level.

Figure S2. Comparative analysis of the Tet(X2)-like orthologs at the amino acid level.

Figure S3. Significance analysis of *tet*(X2) distributions in Europe, Asia and America.

Figure S4. Distribution of *tet*(X2)-like orthologs among different bacterial genus, countries and age groups from microbial genomic bins of human-gut origin.

Figure S5. Comparative analysis of the *tet*(X2) genomic context among *E. coli*, *R. anatipestifer*, *Phocaeicola vulgatus* and *Odoribacter laneus*. Arrows indicate the directions of transcription of the genes, and different genes are shown in different colors. Regions of ≥ 99.0% nucleotide sequence identity are shaded light grey. The *Δ* symbol indicates a truncated gene. IS, insertion sequence. See Table S11 for genomic Type XVIII – XXII definitions.

Table S1. A total of 12,829 non-duplicate metagenomic samples that derived from previous studies (see PMID of Publication).

Table S2. A total of 202,265 MGBs that was constructed from the 12,829 non-duplicate metagenomic samples.

Table S3. The *tet*(X2)-like and non *tet*(X2) orthologs designed in current study.

Table S4. Minimum inhibitory concentration of the *tet*(X)s from metagenomic analysis. TET: tetracycline; DOX: doxycycline; MIN; minocycline; TIG: tigecycline; ERA: eravacycline; OMA: omadacycline.

Table S5. Positive rates of *tet*(X) gene in 31 countries.

Table S6. Positive rates of *tet*(X) carrying MGBs annotated at family level.

Table S7. Positive rates of *tet*(X) carrying MGBs annotated at species level. Table S8. Detail information of the 322 *tet*(X) carrying MGBs.

Table S9. Detail information of the 896 *tet*(X) carrying bacterial isolates.

Table S10. Clusters of the 1218 *tet*(X) positive contigs from the MGBs and bacterial isolates.

Table S11. Detail information of the *tet*(X) genomic context types I to XXII.

